# Contractile force of transplanted cardiomyocytes contributes to heart function after injury

**DOI:** 10.1101/2021.11.23.469715

**Authors:** Tim Stüdemann, Judith Rössinger, Christoph Manthey, Birgit Geertz, Rajiven Srikantharajah, Constantin von Bibra, Aya Shibamiya, Maria Köhne, Antonius Wiehler, J. Simon Wiegert, Thomas Eschenhagen, Florian Weinberger

**Author notes:** Corresponding authors: Thomas Eschenhagen, MD, Department of Experimental Pharmacology and Toxicology, University Medical Center Hamburg-Eppendorf, Germany. Martinistr. 52 20246 Hamburg. Florian Weinberger, MD, Department of Experimental Pharmacology and Toxicology, University Medical Center Hamburg-Eppendorf, Germany. Martinistr. 52, 20246 Hamburg.

## Abstract

Transplantation of pluripotent stem cell-derived cardiomyocytes represents an innovative therapeutic strategy for heart failure. Studies in small and large animals have demonstrated functional recovery of left ventricular function after cardiomyocyte transplantation^1–4^, and first clinical studies are currently underway^5^. Yet, the mechanism of action underlying graft-induced benefit is unknown^6^. Here we demonstrate that transplanted cardiomyocytes actively contribute to heart function. We transplanted cardiomyocytes with an optogenetic off-on switch in a guinea pig cardiac injury model. Light-induced inhibition of engrafted cardiomyocyte contractility resulted in a rapid decrease of left ventricular function that was fully reversible with the offset of photostimulation. Hence, our optogenetic approach demonstrated that transplanted cardiomyocytes actively participate in heart function, supporting the hypothesis that the delivery of new force-generating myocardium can serve as a regenerative therapeutic strategy.

## Main

Ischemic heart disease is the main cause of death globally^7^. The adult heart has minimal regenerative capacity. Myocardial infarction results in a permanent loss of contractile myocardium and regularly leads to the development of heart failure^6,8^. Transplantation of pluripotent stem cell-derived cardiomyocytes represents a regenerative therapeutic concept with great potential for the treatment of heart failure patients^9^. It was successfully applied in various preclinical studies^1–4,10^ and is currently evaluated in the first clinical trials^5^ (e.g. BioVAT-HF, NCT04396899). The goal of this strategy is conceptually different from all current clinically established approaches as it aims at remuscularization, providing new force-developing myocardium to the injured heart. It is intuitive to assume that new myocytes actively participate in the mechanical work of the heart and support its compromised pump function. However, improvement in left ventricular function was also reported in studies with only minimal^11^ or even without cell engraftment^12^, suggesting that other mechanisms are at work here. A recent consensus paper concluded that transplanted cardiomyocytes might contribute to systolic force generation, but formal proof that engrafted cells actively support the mechanical work of the heart is lacking^6^.

Here we generated cardiomyocyte lines with an off-on switch to dissect the mechanism by which cardiomyocyte transplantation improves left ventricular function. We engineered induced pluripotent stem cells with CRISPR/Cas9 to silence cardiomyocyte contractility reversibly. To this end, we knocked-in an inhibitory Pharmacologically Selective Actuator Modul (PSAM^L141F,Y115F^-GlyR, consisting of the chloride-selective ion pore domain of the glycine receptor and mutated ligand-binding domain of the α7 nicotinic acetylcholine receptor) and cytosolic enhanced green fluorescent protein (EGFP)^13^. In a second approach, we used an optogenetic strategy. As light delivery is challenging in the heart, we knocked in an inhibitory luminopsin 4 (iLMO4, consisting of the Gaussia luciferase M23, the improved chloride-conducting channelrhodopsin (iChloC), and enhanced yellow fluorescent protein (EYFP))^14–16^. iLMO4 can be activated by light but also by the luciferase substrate coelenterazine (CTZ). Both constructs were knocked into the AAVS1 safe harbor locus of human induced pluripotent stem cells (iPSC line UKEi001-A; Supplementary Figure 1)^17^. PSAM-GlyR, iLMO4 iPSCs, as well as iPSC from the parental line UKEi001-A (WT) were differentiated into cardiomyocytes (average troponin T positivity: 71% (PSAM-GlyR); 84% (iLMO4); 95% (WT); Figure 1a and Supplementary Figure 2). Transgene expression in PSAM-GlyR and iLMO4 cardiomyocytes was >95% (Supplementary Figure 2b and d).

**Figure 1:**
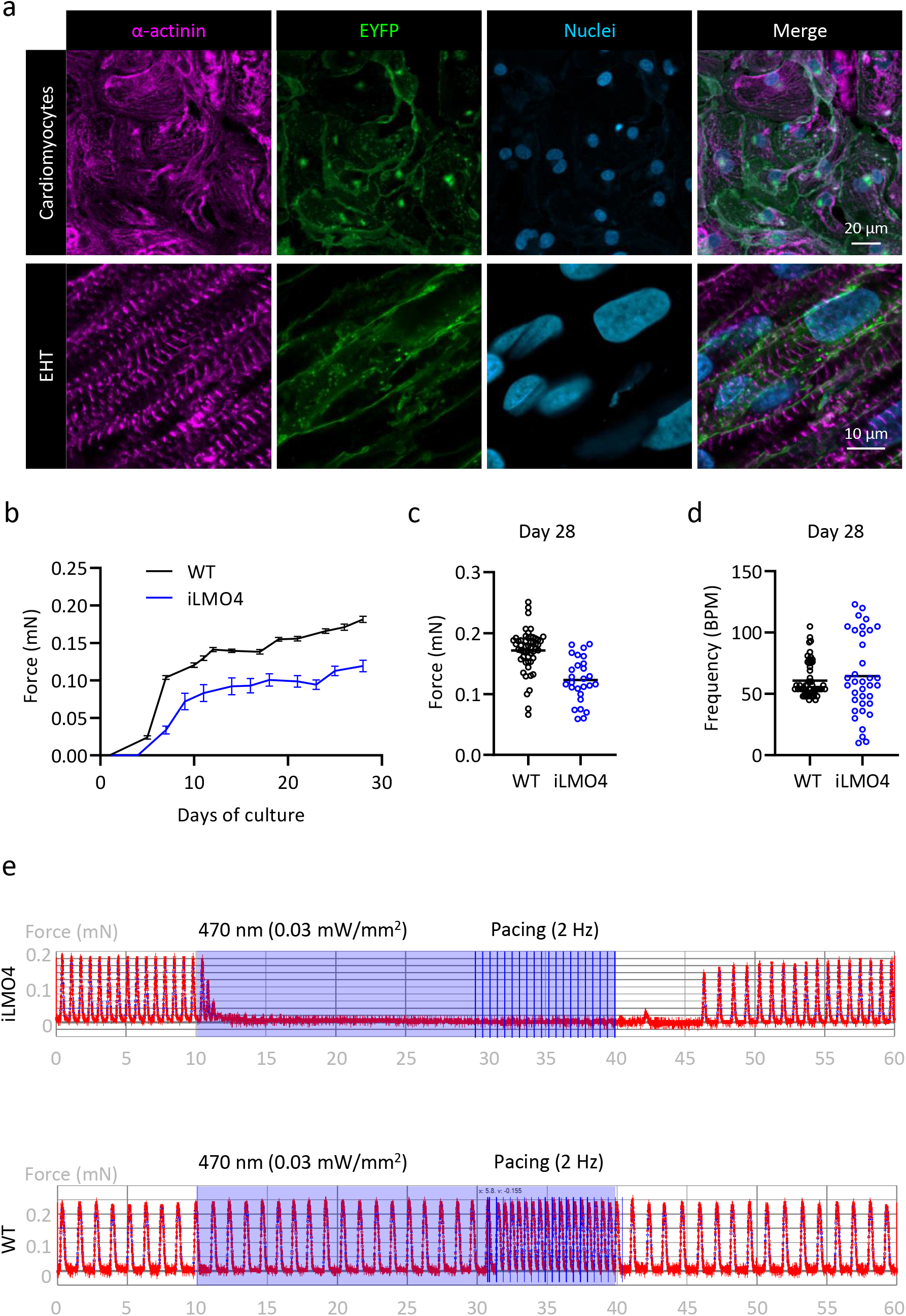
Photostimulation reversibly stops iLMO4-cardiomyocyte contractility. **a**) Immunohistology of iLMO4 cardiomyocytes and an iLMO4 EHT, respectively. **b** and **c**) Physiological characterization of iLMO4-EHTs. **b)** Force development of WT and iLMO4 EHTs over time. Data from three different batches per cell line (4-41 EHTs per batch) were averaged. Mean ± SEM are shown. **c**) Force and **d**) frequency analysis after four weeks in culture from three different batches (n=4-41 EHTs per batch and cell line, each data point represents one EHT). **e)** Original force recording under photostimulation from iLMO4 (top) and WT EHTs (bottom). Vertical blue lines indicate electrical pacing (2 Hz; 2 V; 4 ms impulse duration).

For functional characterization, engineered heart tissue (EHT) was generated from PSAM-GlyR and iLMO4 cardiomyocytes^18^ (Figure 1a and Supplementary Figure 3a). PSAM-GlyR and iLMO4 EHTs developed similar to WT EHTs and started to beat coherently after 7-10 days in culture. Force and frequency of PSAM-GlyR EHTs were similar to WT EHTs (Supplementary Figure 3b). iLMO4 EHTs showed greater variability in force and frequency than WT EHTs (average force 0.12±0.01 mN vs. 0.17±0.01 mN in WT; Figure 1b-d). PSAM-GlyR EHTs were exposed to a chimeric ion channel agonist (Pharmacologically selective effector module, PSEM^89S^) to test the off-on switch at increasing concentrations. PSEM^89S^ (30-100 μM) stopped PSAM-GlyR EHT contractility. This effect was completely reversible after wash-out (Supplementary Figure 3d and e) and did not occur in WT EHTs (Supplementary Figure 3f).

iLMO4 EHTs were intermittently exposed to light pulses (30 s, 470 nm). Photostimulation had no effect in WT EHTs, but induced an immediate stop of contractility in iLMO4 EHTs, which resumed 5-10 seconds after photostimulation (Figure 1e and Supplementary Figure 4 and Movie 1). Alternatively, iLMO4 can be activated by CTZ. Application of CTZ resulted in light emission (Supplementary Figure 4b) and stopped EHT contractility. However, this strategy had several limitations: i) high CTZ concentration was needed (~300 μM), ii) the effect was inconsistent, iii) the off- and on switch kinetics were slow (off switch 5-20 min; on-switch 20-60 min after wash-out) and iv) not all EHTs fully recovered after CTZ wash-out (Supplementary Figure 4c). Therefore, we focused on direct photostimulation to manipulate iLMO4 cardiomyocyte contractility.

To assess whether the contractile force of transplanted cardiomyocytes participates in left ventricular function, PSAM-GlyR, iLMO4, or WT cardiomyocytes were injected transepicardially in a subacute guinea pig injury model (20×10^6^ cardiomyocytes per animal in pro-survival cocktail, n=30; Supplementary Figure 5a), hypothesizing that switching off contractility in engrafted cardiomyocytes would result in a decrease in left ventricular function. Initially, we focused on PSAM-GlyR cardiomyocytes (Supplementary Figure 6a-c) because the contractile function of PSAM-GlyR EHTs more closely mirrored that of WT EHTs.

However, whereas PSEM^89S^ did not affect force in WT EHTs in vitro, infusion of PSEM^89S^ exerted a profound negative chronotropic and inotropic effect in Langendorff-perfused guinea pig hearts, irrespective of cardiomyocyte transplantation (Supplementary Figure 6d and e). This effect might be caused by PSEM^89S^-mediated off-target stimulation of muscarinergic or nicotinic acetylcholine receptors at the sinoatrial node or parasympathetic ganglia, respectively. Electrical pacing reversed the negative chronotropic effect, but the negative inotropic response persisted and could not be abolished with a nicotinic (hexamethonium) or a muscarinergic (atropine) antagonist. Therefore, we switched to the optogenetic approach because the non-selective negative inotropic PSEM89S effect might have masked effects of turning off contractility in the human grafts.

Hearts were harvested 28 days after transplantation of iLMO4 or WT cardiomyocytes (Figure 2a). Hearts showed transmural scarring. Scar size was slightly larger in hearts that had received WT cardiomyocytes (27±2% WT vs. 20±1% iLMO4, Figure 2a). Human cardiomyocytes partially remuscularized the scar (9±2% of scar size in WT vs. 13±3% in iLMO4; Figure 2b). Human origin was identified by EYFP immunostaining (Figure 2c-e). The engrafted cardiomyocytes almost exclusively expressed the ventricular isoform of the myosin light chain (Figure 2f). Still, they showed signs of immaturity compared to host cardiomyocytes (lower sarcomeric organization, peripheral myofibrils, smaller cell size, circumferential connexin43, and cadherin expression, Figure 2e-h). The grafts showed extensive zones of close proximity with host myocardium (Figure 2d), and human cardiomyocytes (EYFP^+^) formed immature N-cadherin^+^ adherens junctions and connexin43^+^ gap junctions with host cardiomyocytes (Figure 2g and h), indicating cell-cell-coupling.

**Figure 2:**
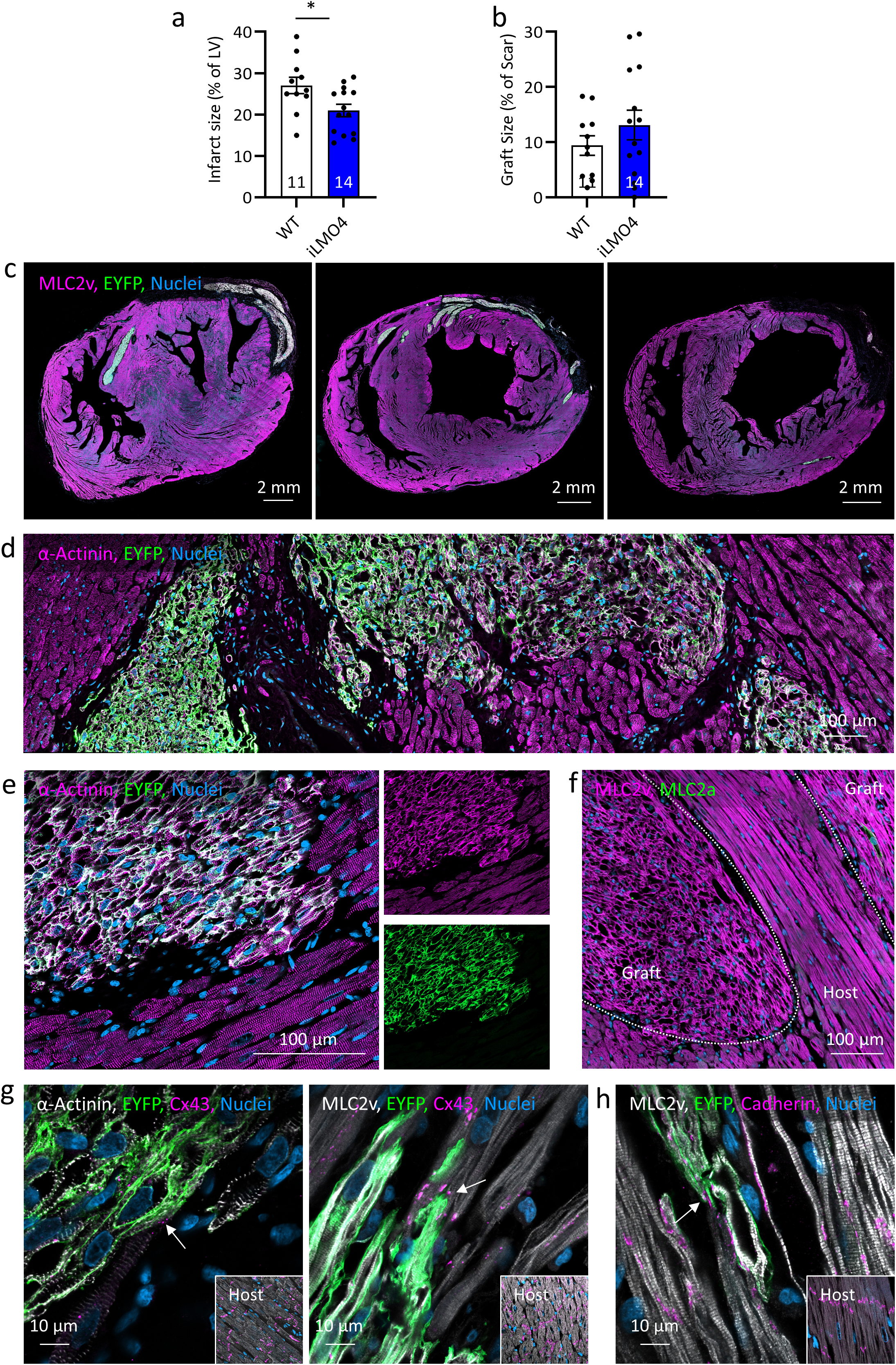
iLM04 transplantation partially remuscularized the injured heart. **a)** Infarct size quantification as percentage of the left ventricle. **b)** Quantification of graft size as percentage of scar area (n=11-14; each data point represents one heart). Mean ± SEM are shown. **c)** Short axis sections from three different hearts four weeks after iLM04-cardiomyocyte transplantation demonstrating the varying degree of remuscularization. **d** and **e)** Graft-host interaction in lower **(d)** and higher magnification **(e). f)** Analysis of myosin light chain isoform expression in the human graft. **g** and **h)** Analysis of cell-cell contacts between engrafted and host cardiomyocytes. **g)** High magnification connexin43 images from 2 different hearts. **h)** High magnification N-cadherin staining. Insets show host tissue stained with the respective antibody combination.

Echocardiography was performed at baseline, before transplantation, and four weeks after transplantation. Cardiomyocyte transplantation stabilized left ventricular function (fractional area change (FAC) 42±1% at baseline, 28±3% (WT) and 32±2% (iLMO4) after injury and 31±2% (WT) and 33±3% (iLMO4) four weeks after transplantation (Figure 3A). We employed ex-vivo Langendorff-perfusion^19^ for an accurate functional analysis instead of in vivo imaging for two reasons: i) in vivo imaging (by echocardiography) has limited precision, particularly after repeated thoracotomies^20^ and ii) switching off contractility of the human grafts was expected to exert only a small effect because electrical coupling between graft and guinea pig host myocardium was shown to be incomplete and spatially and temporally heterogeneous^21^. In contrast to the chemogenetic strategy, photostimulation did not affect the guinea pig heart function. After a run-in period (> 15 minutes, during which LV-pressure amplitude increased by 15.1±1.8 mmHg), repetitive (blue light pulses (470 nm, 30-60 seconds) were applied to the anterior epicardial surface of spontaneously beating guinea pig hearts (average beating frequency 161±6 bpm; Supplementary Figure 7a and Movie 3). Photostimulation resulted in an instantaneous drop in left ventricular pressure in 5 out of 10 hearts with iLMO4 cardiomyocytes (pressure amplitude prior light application (baseline): 60.4±3.8 mmHg (n=10); during light application: 59.9±3.7 mmHg (n=10); ∆LVDP -0.5±0.3 mmHg (n=10); relative ∆LVDP between baseline and photostimulation -0,8%). The maximal effect was a drop of 2.6 mmHg accounting for 4.7% (Figure 3b-e and Supplementary Figure 7b). The effect size fits well to the calculated graft mass (~40 mg; calculated graft mass= left ventricular mass [2.3 g (average heart mass) x 0.6 x 0.20 (scar size; % of left ventricle)] x 0.13 (graft size % of scar size)) that accounts for ~2-3% of the left ventricular mass (~1.5 g, left ventricular mass=2.3 g (average heart mass) x 0.6) The effect of blue light reversed within a few seconds (~5-10 s) after the termination of the light pulse (relative ∆LVDP between photostimulation and recovery +1,5%). In contrast, no heart that had received WT cardiomyocytes (n=5) reacted to blue light exposure (pressure amplitude prior light application (baseline): 59.5±4.2 mmHg (n=5); during light application: 59.7±4.2 mmHg (n=5); ∆LVDP +0.4±0.1 mmHg (n=5); relative ∆LVDP between baseline and photostimulation +0,8%; relative ∆LVDP between photostimulation and recovery -0.2%; Figure 3a and b and Supplementary Figure 7b). Photostimulation with red light (660 nm) was used as an internal control and did not affect left ventricular function in iLMO4 or WT transplanted hearts (Figure 3b, c and e, and Supplementary Figure 7b), excluding non-specific effects (e.g. temperature effects). As CTZ had sustained detrimental effects on EHTs, we only applied it at the end of the Langendorff experiment to verify cardiomyocyte viability. CTZ application demonstrated that the transplanted cardiomyocytes were viable and that the graft was functionally vascularized (Figure 3f).

**Figure 3:**
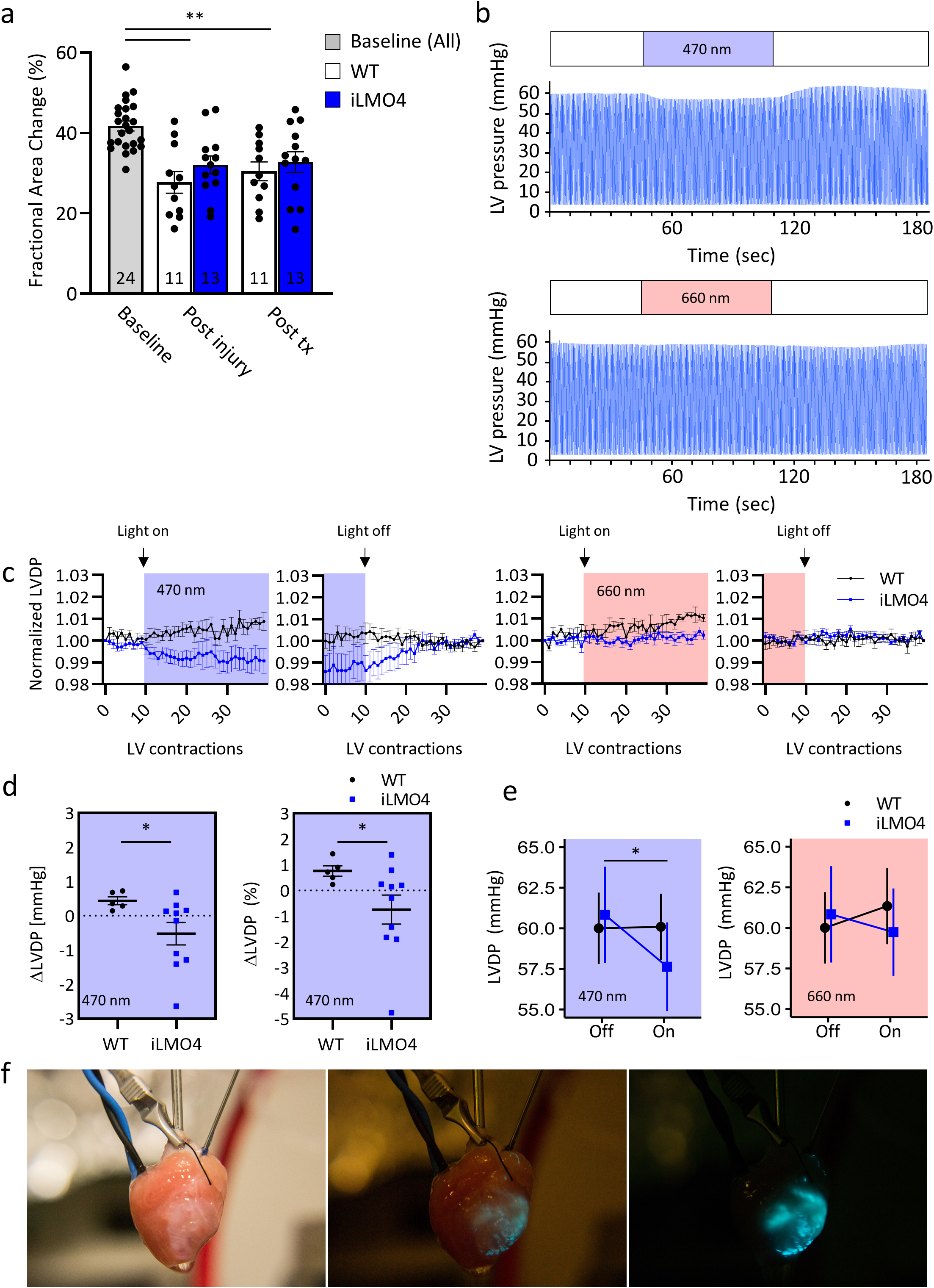
Engrafted cardiomyocytes actively participate in left ventricular function. **a)** Serial echocardiographic analysis of heart function. Fractional area change values for baseline, post-injury and 4 weeks after transplantation (n=11 (WT) and n=13 (iLMO4)). Statistical analysis was determined by two-way ANOVA followed by Tukey’s-Test for multiple comparisons. **P<0.01. **b)** Original left ventricular pressure recording from an iLMO4 heart during photostimulation with blue (470 nm) and red light (660 nm) respectively. **c)** Normalized LVDP from WT (n=5) and iLMO4 (n=10) hearts. Shown are ten beats prior to photostimulation, and the first 30 beats during photostimulation (left), and the last ten beats during photostimulation followed by the first 30 beats after photostimulation (right). LVDP was normalized to the first depicted beat or the least depicted beat. **d)** Absolute (left) and relative (right) difference between LVDP between baseline (prior to photostimulation) and photostimulation with blue light. The first five beats (0-5 from **c)** prior to photostimulation and five beats during photostimulation (35-40 from **c)** were averaged. **e)** Comparison of all heart beats during photostimulation and baseline. A mixed model was used for statistical analysis. **f)** Photographs of an iLMO4 heart after the application of CTZ (~300 μM). LVDP: left ventricular developed pressure.

Preclinical models have repeatedly demonstrated the efficacy of cardiomyocyte transplantation to stabilize or even improve left ventricular function. Yet, so far, the mechanism by which the transplanted cells exert their effect was unknown. Our study now provides evidence that transplanted cardiomyocytes actively support the contractile heart function, fulfilling a central criterion of cardiac remuscularization. The overall effect on heart function was small. Yet, there are several reasons to conclude that it is a true effect: i) Hearts that received WT cardiomyocytes did not react to blue light exposure, ii) the internal control with red light did not result in a drop of left ventricular function, iii) the effect size fits well when comparing the engrafted myocardial mass to the left ventricular myocardial mass (40 mg to 1.5 g).

Additionally, i) differences between the guinea pig and human cardiac physiology, ii) incomplete electrical coupling^21^, iii) the immaturity of the transplanted cells, iv) the unstructured myocyte orientation, and v) the myocardial edema, particularly in the scarred region, in the Langendorff-system^22^ might have attenuated the extent of the force that contributes to left ventricular function in this model. There is evidence that coupling is more efficient in a large animal model^23^ and that the cells mature over time^23^, indicating that the beneficial contractile effects after transplantation in humans can be larger.

Overall, our findings are encouraging regarding ongoing and pending regenerative clinical trials for heart failure patients, as they demonstrate that reconstitution of lost myocardium can serve as a conceptually new strategy to improve heart function.

## Methods

### Generation of the PSAM-GlyR and iLMO4 iPSC-line

AAVS1-CAG-hrGFP was a gift from Su-Chun Zhang (Addgene plasmid # 52344 ; http://n2t.net/addgene:52344 ; RRID:Addgene_52344)^17^. CAG::PSAML141F,Y115F:GlyR-IRES-GFP was a gift from Scott Sternson (Addgene plasmid # 32480 ; http://n2t.net/addgene:32480 ; RRID:Addgene_32480)^13^. iLMO4 plasmids were a gift from Ute Hochgeschwender (Central Michigan University, USA)^15^. The plasmid vectors contained PSAM-GlyR or iLMO4 under the control of a CAG promoter, homology arms (~800 bp in size each) for the AAVS1 locus, and a puromycin resistance cassette upstream of the promoter. PSAM-GlyR was linked to EGFP by a 2A linker. Nucleofection with the Cas9 RNP was conducted using a 4D-Nucleofector (Lonza) according to the manufacturer's protocol. The selected sgRNA targeted the AAVS1 between Exon 1 and 2 (Supplementary Figure 1). Positively edited cells were enriched by FACS (BD Aria Fusion) and seeded as single cells.

### Genotyping

Correct transgene integration was verified by Southern blotting. EcoRI was used to digest genomic DNA. Transgene integration introduced two new EcoRI cutting sites. A 5' probe allowed detection of a 4.9 kb fragment in case of transgene integration or detection of an 8.3 kb fragment on the control allele. Additionally, an internal probe was used to detect a 6.8 kb fragment only on successfully targeted alleles. Moreover, PCR amplification followed by Sanger sequencing was used to verify correct transgene integration.

### Human iPSC culture and cardiac differentiation

UKEi001-A was reprogrammed with a Sendai Virus (CytoTune iPS Sendai Reprogramming Kit, ThermoFisher). Expansion and cardiac differentiation of iPSC were performed as recently described^1^. In brief, iPSCs were expanded in FTDA medium on Geltrex-coated cell culture vessels. Formation of embryoid bodies was performed in spinner flasks, followed by differentiation in Pluronic F-127-coated cell culture vessels with a sequential administration of growth factor- and small-molecule-based cocktails to induce mesodermal progenitors, cardiac progenitors, and cardiomyocytes. Dissociation of differentiated cardiomyocytes was performed with collagenase. Cardiomyocytes designated for transplantation underwent heat shock 24 hours before dissociation as previously described ^3,23,24^ and were cryopreserved. Antibodies used for characterization are listed in Table 1.

**Table 1.** Primary antibodies for flow cytometry.

| Antigen | Supplier | Titer |
| --- | --- | --- |
| **SSEA3 (AlexaFluor® 647)** | BD Pharmingen, Clone MC-63, Cat. 561145 | 1:50 |
| **Isotype SSEA3** | BD Pharmingen, clone MOPC/21, Cat. 557714 | 1:50 |
| **Cardiac troponin T (APC)** | Miltenyi Biotec, clone REA400, Cat. 130-120-403 | 1:50 |
| **Isotype cardiac troponin T** | Miltenyi Biotec, clone REA293, Cat. 130-120-709 | 1:50 |

### Flow cytometry

Single-cell suspensions of iPSCs were blocked with 5% FBS in PBS and stained. Single-cell suspensions of iPSC-derived cardiomyocytes were fixated in Histofix (Roth A146.3) for 20 min at 4 °C and transferred to FACS buffer, containing 5% FBS, 0.5% Saponin, 0.05% sodium azide in PBS. Antibodies are listed in Table 1. Samples were analyzed with a BD FACSCanto II Flow Cytometer and the BD FACSDiva Software 6.0 or BD FlowJo V10. An example of the gating strategy is given in Supplementary Figure 5b.

### EHT generation and analysis

EHTs were generated from cells and fibrinogen/thrombin as previously described^1^. In brief, spontaneously beating WT or iLMO4 cardiomyocytes were digested with collagenase II (Worthington, LS004176; 200 U/ml Ca^2+^-free HBSS (Gibco, 14175-053) with 1 mM HEPES (pH 7.4), 10 μM Y-27632, and 30 μM N-benzyl-p-toluene sulfonamide (TCI, B3082)) for 3.5 hr at 37°C (5% CO_2_, 21% O_2_). The dissociated cells were resuspended in Ca^2+^-containing DMEM with 1% penicillin/streptomycin. Cell concentration was adjusted to 10–15×10^6^ cells/ml. Fibrin-based human EHTs were generated in agarose casting molds with solid silicone racks^25^ (100 μl per EHT, 1×10^6^ cells). The culture medium was changed on Mondays, Wednesdays, and Fridays. After ~7 days in culture, human EHTs displayed spontaneous coherent, regular beating deflecting the silicone posts that allowed video-optical contraction analysis. Force measurement and photostimulation of iLMO4 EHTs were performed as described by Lemme et al.^26^. In brief, analysis of contractile force was performed by video-optical recording on a setup available from EHT Technologies. The contraction peak analysis was performed during spontaneous beating under red light illumination. Electrical pacing was performed with pacing electrodes for 24 well plates (EHT Technologies).

### Animal care and experimental protocol approval

The investigation conforms to the guide for the care and use of laboratory animals published by the NIH (Publication No. 85-23, revised 1985) and was approved by the local authorities (Behörde für Gesundheit und Verbraucherschutz, Freie und Hansestadt Hamburg: N098/2019).

### Injury model and cardiomyocyte transplantation

Myocardial injury was induced as previously described^1^. Cryoinjury of the left ventricular wall was induced in female guinea pigs (300-470 g, 8-9 weeks of age, Envigo). Cardiomyocyte transplantation was performed during a repeated thoracotomy seven days after injury. 20×10^6^ cardiomyocytes, resuspended in pro-survival cocktail^2,4^ (Matrigel (~50% v/v), cyclosporine A (200 nM, pinacidil (50 μM), IGF-1 (100 ng/ml) and Bcl-X_L_ BH4 (50nM), total volume: 150 μl) were injected into three separate injection sites, e.g., one injection in the central lesion and two injections in the flanking lateral border zones. Surgeons were blinded regarding the injected cell line. Guinea pigs were immunosuppressed with cyclosporine beginning three days prior to transplantation (7.5 mg/kg body weight/day for the first three postoperative days and 5 mg/body weight/day for the following 25 days; mean plasma concentration: 435 μg/l) and methylprednisolone (2 mg/kg body weight/day).

### Echocardiography

Echocardiography was performed with a Vevo 3100 System (Fujifilm VisualSonics) as previously described^1^. Analysis was performed in a blinded manner. One animal showed no decrease in fractional area change one week after injury and was therefore excluded from further analysis.

### Histology

Hearts were sectioned in four to five slices after fixation in Histofix. Serial paraffin sections were acquired from each slice and used for histology and immunohistology. Antigen retrieval and antibody dilution combinations used are summarized in Table 2. The primary antibody was either visualized with the multimer-technology based UltraView Universal DAB Detection kit (Ventana® BenchMark® XT; Roche) or a fluorochrome-labeled secondary antibody (Alexa-conjugated, Thermofisher). Confocal images were acquired with an LSM 800 (Zeiss). Whole short-axis sections were acquired with stitched images in 20x magnification using the LSM 800 (Zeiss). For morphometry, images of dystrophin stained sections were acquired with a Hamamatsu Nanozoomer whole slide scanner and viewed with NDP software (NDP.view 2.6.13). Infarct size was determined with a length-based approach as described previously^10^. Graft size was measured in dystrophin-stained short-axis sections and expressed as a percent of the scar area measured in the same section using the NPD2.view-software.

**Table 2.** Primary antibodies for histology.

| Antigen | Supplier | Antigen retrieval | Titer |
| --- | --- | --- | --- |
| **MLC2A** | BD Pharmingen, clone S58-205, Cat. 565496 | Citrate buffer | 1:500 |
| **MLC2V** | Proteintech, pAb, Cat.10906-1-AP | Citrate buffer | 1:250 |
| **Alpha-Actinin** | Sigma-Aldrich, clone EA-53<br>Cat. A7811 | Proteinase K | 1:600 |
| **Dystrophin** | Millipore, clone 1808, Cat. MAB1645 | EDTA | 1:150 |
| **Ku80** | Cell Signaling Technology, clone C48E7,<br>Cat. CST-2180 | Citrate buffer | 1:800 |
| **Connexin 43** | BD Transduction Laboratories, clone<br>2/Connexin-43, Cat. 610062 | Citrate buffer | 1:400 |
| **Connexin 43** | Abcam, pAb, Cat. ab11370 | Citrate<br>buffer/Proteinase K | 1:400 |
| **N-Cadherin** | Sigma-Aldrich, clone CH19, Cat. C1821 | Citrate buffer | 1:400 |
| **GFP** | Abcam, clone 3H9, Cat. ab252881 | Citrate<br>buffer/Proteinase K | 1:500 |
| **GFP** | Abcam, pAb, Cat. ab6556 | Proteinase K | 1:500 |

### Langendorff-perfusion

Guinea pigs (bodyweight 430–570 g) were injected with heparin (1000 U/kg, s.c.) and anesthetized with midazolam (1 mg/kg mg/kg, i.m.), medetomidine (0,2 mg/kg mg/kg, i.m.) and fentanyl (0,02 mg/kg mg/kg, i.m.). Hearts were excised and immediately immersed in ice-cold modified Krebs–Henseleit solution containing (in mM) NaCl 120, KCl 4.7, MgSO_4_ 1.2, NaHCO_3_ 25.0, KH_2_PO_4_ 1.2, glucose 11.1, Na-pyruvate 2.0, and CaCl_2_ 1.8. Lidocaine (170 μM) was added to prevent premature ventricular contractions. The aorta was quickly cannulated, and the heart was connected to a custom-made Langendorff apparatus. The heart was retrogradely perfused using hydrostatic pressure (~55 mmHg) with warm (36.5±1 °C) modified Krebs-Henseleit solution equilibrated with a mixture of 95% O_2_ and 5% CO_2_ (pH 7.35–7.40). Left ventricular pressure was recorded with a self-assembled balloon catheter, and data were acquired using Chart5 (ADInstruments). After a stabilization period of (>15 min), hearts were photostimulated by continuously illuminating the anterior wall of the left ventricle (peak excitation 470 nm, (maximal light intensity 712 mW pE4000, coolLED) for 30-60 seconds. At least three photostimulations with blue light were performed per heart, separated by 1 min each. The photostimulation periods with blue light were followed by a photostimulation with red light (660 nm, maximal light intensity 570 mW) for 30-60 seconds. After the termination of the photostimulation procedures, CTZ (~ 300 μM) was applied in a subset of hearts (n=3).

### Statistics

Statistical analyses were performed with GraphPad Prism 9, USA and R, Vienna, Austria. Comparison among two groups was made by two-tailed unpaired Student's t-test. To physiologically compare EHT from either PSAM-GlyR or iLMO4 cardiomyocytes with WT EHTs nested two-tailed unpaired Student's t-test was used. One-way ANOVA followed by Tukey's-Test for multiple comparisons were used for more than two groups. When two factors affected the result (e.g. timepoint and group), 2-way ANOVA analyses and Tukey's-Test for multiple comparisons were performed. Error bars indicate SEM. P-values are displayed graphically as follows: * p < 0.05, ** p < 0.01, *** p < 0.001. Additionally, statistical analyses to assess the effect of photostimulation were performed in R with linear mixed models (packages “lmer”, “sjPlot”, “emmeans”). We regressed LVDP against the following predictors: Time, light (off-blue-red, dummy coded in two variables) and cell type (WT-iLMO4) and interactions of light and cell type. Intercepts and all within-animal factors (e.g. time) were estimated on the animal level. The between-animal factor cell typeas estimated on the group level. We z-scored all continuous predictors per animal

## Author contributions

T.S. performed experiments (generation of the PSAM-GlyR and iLMO4 cell lines, iPSC characterization, cardiac differentiation, animal procedures, performed echocardiography, histology, and Langendorff-experiments), analyzed data, and prepared the manuscript. J.R. and C.M. performed EHT experiments. B.G. conducted animal procedures, performed and analyzed echocardiography, and performed Langendorff-experiments. R.S. performed histological studies. C.v.B. conducted animal procedures, performed and analyzed echocardiography. A.S. and M.K. performed experiments (iPSC cell culture, cardiac differentiation). A.W. performed statistical analysis. J.S.W. provided technical support and conceptual advice for optogenetic experiments. T.E. designed the project, acquired funding, supervised experiments, and prepared the manuscript. F.W. designed the project, supervised and performed experiments (animal procedures, echocardiography, histology, and Langendorff-experiments), acquired funding, and prepared the manuscript.

## Acknowledgments

We thank Ute Hochgeschwender, M.D. (Central Michigan University) for providing luminopsin plasmids. We thank Kristin Hartmann (UKE, Mouse Pathology Core Facility) for technical assistance in immunohistochemistry and Dr. Ingke Braren (UKE, Vector Facility) for help with the cloning strategies for PSAM-GlyR and iLMO4 knock-in. We would like to thank Dr. Irm Herrmans-Borgmeyer (Center for Molecular Neurobiology Hamburg, UKE) for help with Southern Blot genotyping and Christiane Pahrmann for help with bioluminescence measurements. We would like to acknowledge Jutta Starbatty, Thomas Schulze, and Birgit Klampe for technical assistance. We would like to appreciate the contribution of Dr. Sandra Laufer during reprogramming the UKEi001-A line. Flow cytometry was conducted in the FACS Core Facility, UKE. We would like to particularly thank the laboratory animal facility staff (UKE) for their support. This work was supported by a Translational Research Grant from the German Centre for Cardiovascular Research (DZHK; 81X2710153 to TE), the European Research Council (ERC-AG IndivuHeart to TE), and the German Research Foundation (DFG; WE5620/3-1 to FW, WI 4485/3-2 to JSW).

## Data Availability

All the data supporting the findings from this study are available within the article and its supplementary information or are available from the corresponding author upon request.

---

**Supplementary Figure 1:**
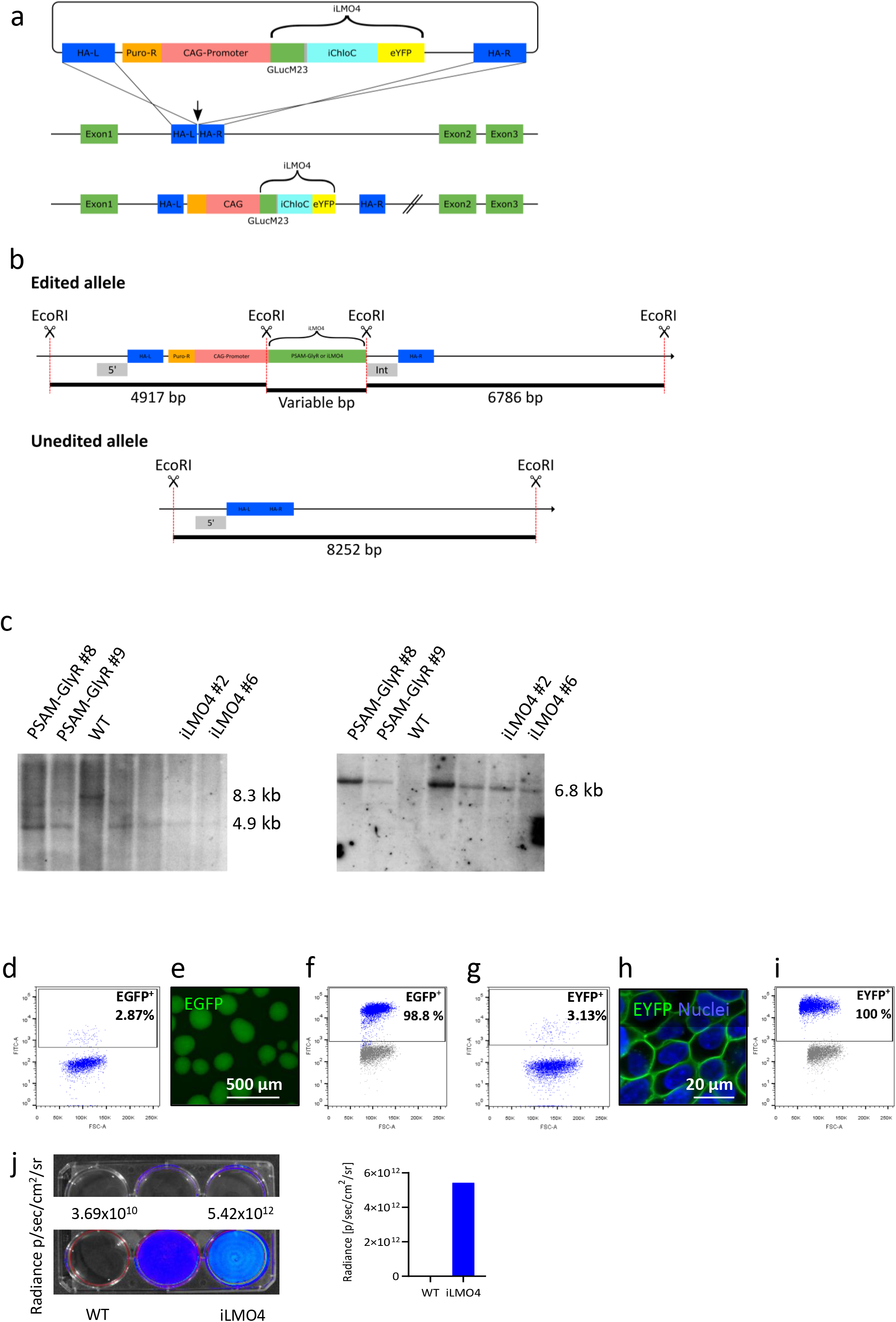
Generation of PSAM-GlyR and iLMO4 iPSC lines. **a)** Knock-in strategy for the iLMO4 iPSC line. PSAM-GlyR knock-in was performed accordingly. **b**) Southern blot genotyping strategy. Two probes (5`and Int (internal) were used to confirm correct transgene integration and homozygosity. **c**) Southern blots from PSAM-GlyR and iLMO4 iPSC-clones with the 5` (left side) and Int-probe (right side). **d)** Flow cytometry of targeted iPSCs (PSAM-GlyR). **e)** Embryoid bodies derived from PSAM-GlyR iPSC after clonal selection. **f)** Flow cytometry from PSAM-GlyR iPSCs. **g**) Flow cytometry of targeted iPSC (iLMO4). **h**) iLMO4 iPSCs after clonal selection. **i**) Flow cytometry from iLMO4 and PSAM GlyR iPSCs. **j**) Bioluminescence measurements in WT and iLMO4 iPSCs.

**Supplementary Figure 2:**
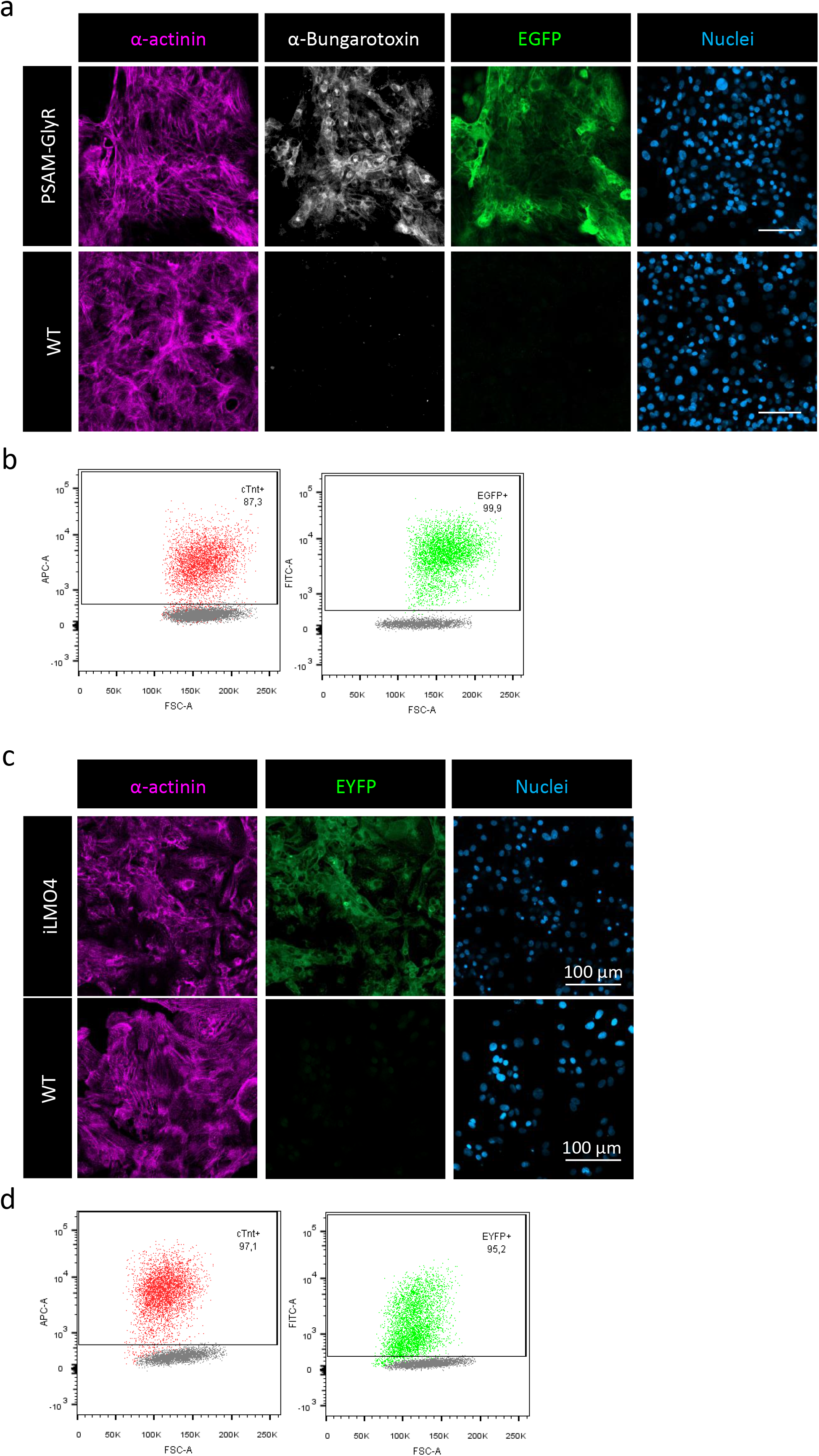
Cardiac differentiation of PSAM-GlyR and iLMO4 iPSCs. **a)** Cardiomyocytes differentiated from PSAM-GlyR iPSCs. α-Bungarotoxin was used to stain the nicotinergic acetylcholine receptor ligand binding domain. **b)** Flow cytometry from PSAM-GlyR cardiomyocytes stained for cardiac troponin t (red) and native EGFP (green). **c**) Cardiomyocytes differentiated from iLMO4 iPSCs. **d**) Flow cytometry from iLMO4 cardiomyocytes stained for troponin t (red) and native EYFP (green).

**Supplementary Figure 3:**
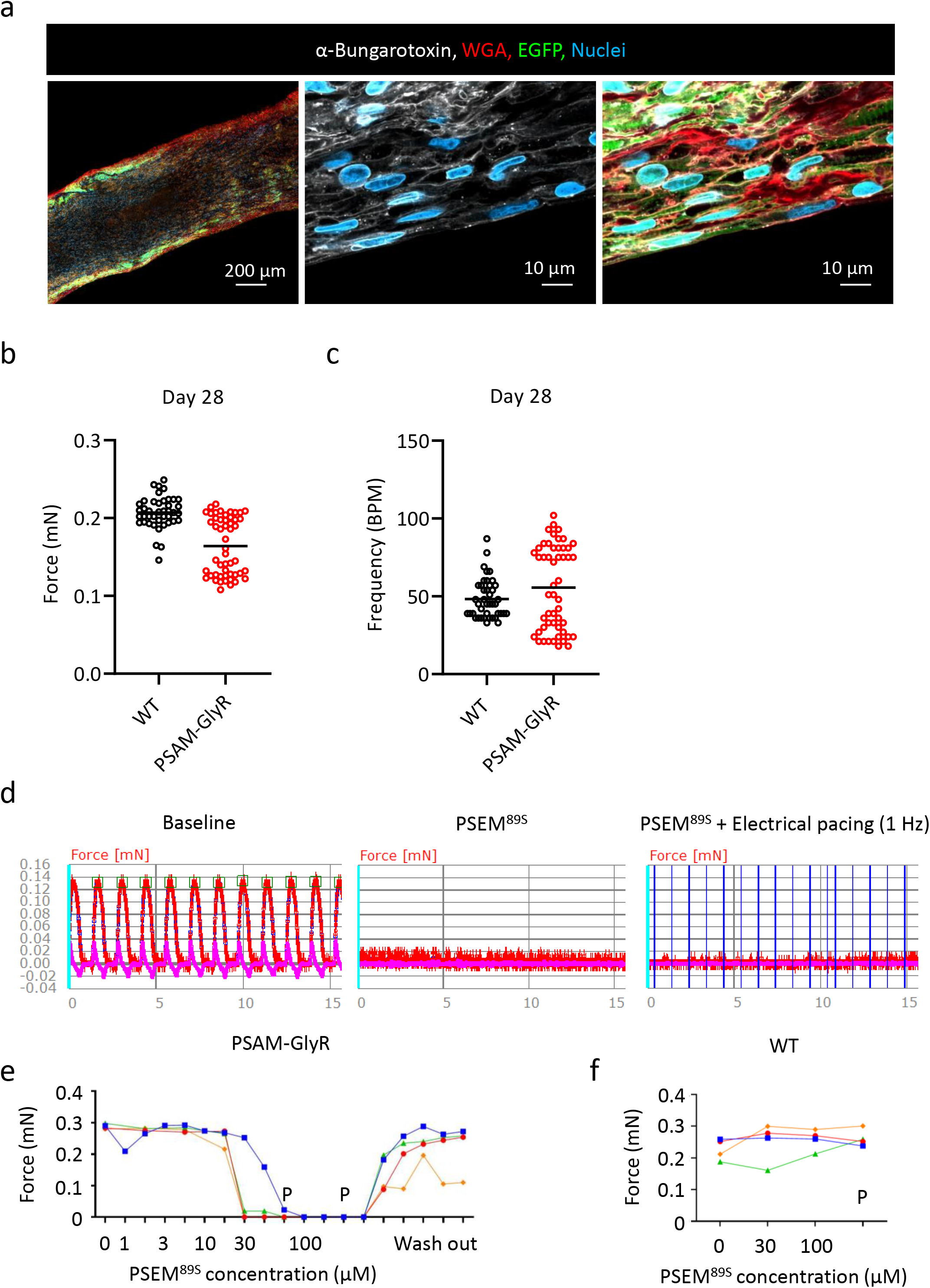
Characterization of PSAM-GlyR EHTs. **a)** Immunofluorescence of a longitudinal PSAM-GlyR EHT section in low and high magnification. α-Bungarotoxin was used to stain the nicotinergic acetylcholine receptor ligand binding domain. **b** and **c**) Physiological PSAM-GlyR EHT characterization. Data for force and frequency between WT and PSAM-GlyR EHTs after one month in culture from three different batches (n=3-21 EHTs per batch and cell line; each dot represents one EHT) are shown. **d**) Original force recording from one PSAM-GlyR EHT under baseline conditions and after incubation with PSEM^89S^ (100 μM) for ten minutes in the absence and presence of external electrical pacing. Vertical blue lines indicate electrical pacing pulses (1 Hz, 2 V, 4 ms impulse duration). **e** and **f)** Force analysis of four different PSAM- GlyR (**e**) and WT EHTs (**f**) under increasing PSEM^89S^ concentrations. P indicates electrical pacing. Measurements were performed one minute and ten minutes after PSEM^89S^ application. n=4, all EHTs shown individually.

**Supplementary Figure 4:**
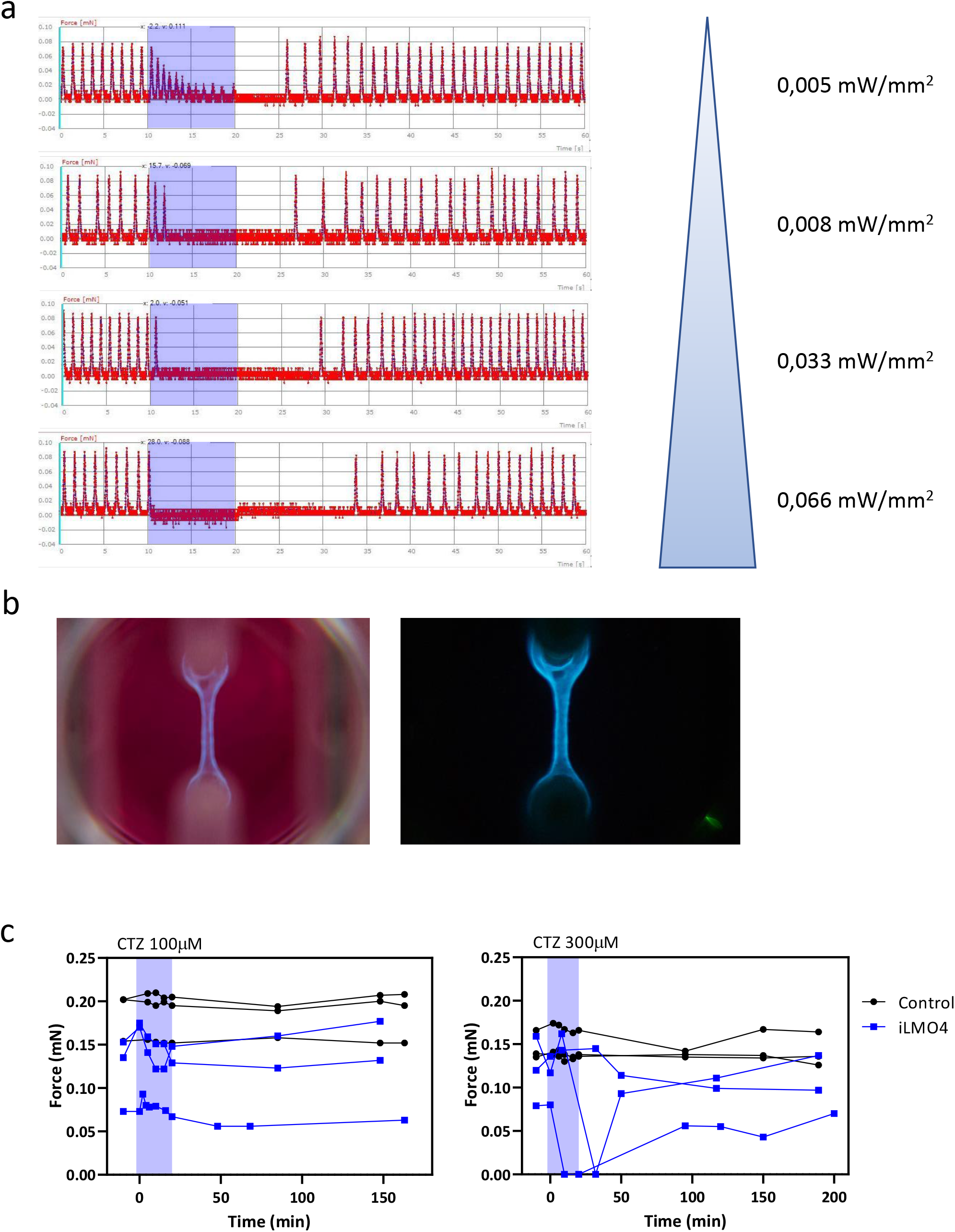
iLMO4 EHT silencing. **a)** Original recordings from iLMO4 EHTs during photostimulation with increasing light intensity. **b**) Photographs of iLMO4 EHTs after incubation with CTZ (300 μM). **c**) CTZ (100 and 300 μM) effect on WT and iLMO4 EHT contractility. Data from three different individual EHTs per group is shown.

**Supplementary Figure 5a:**
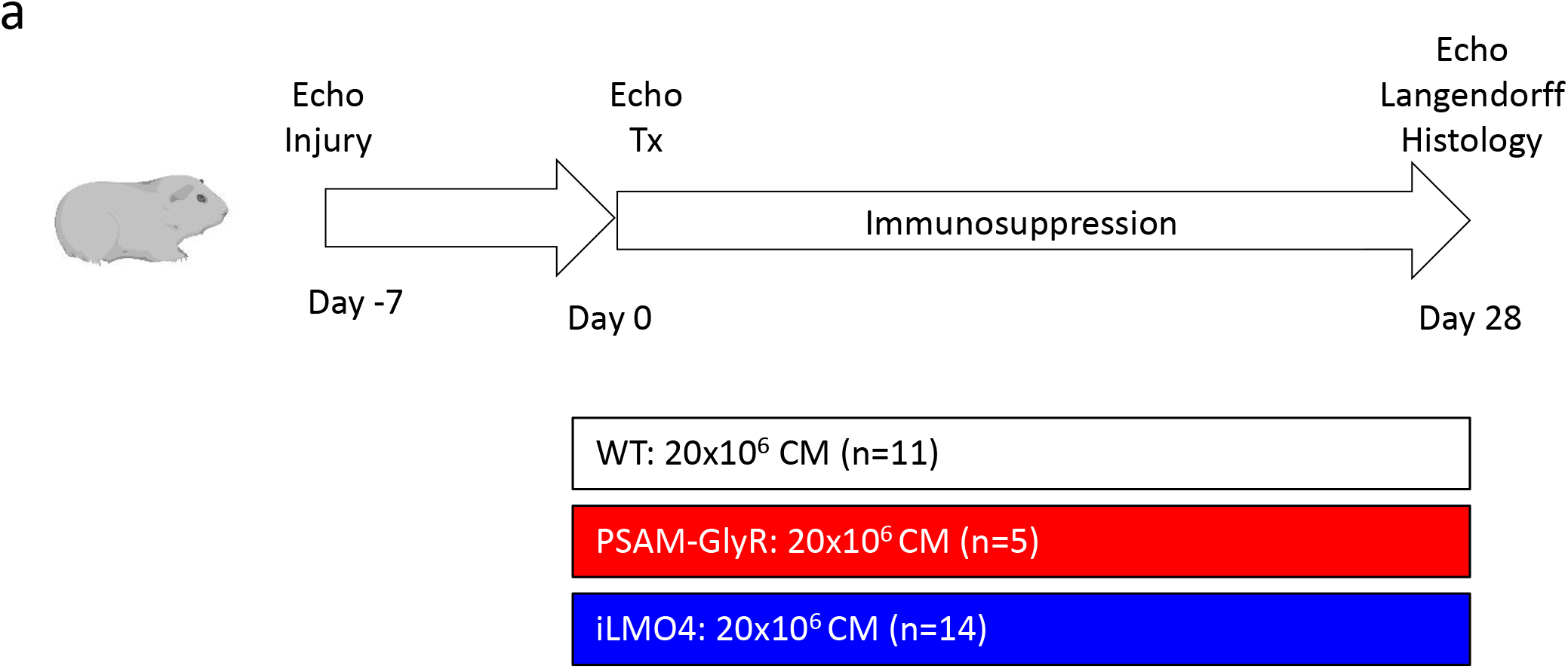
Study protocol. Created with BioRender.com (EH238FM0NG)

**Supplementary Figure 5b:**
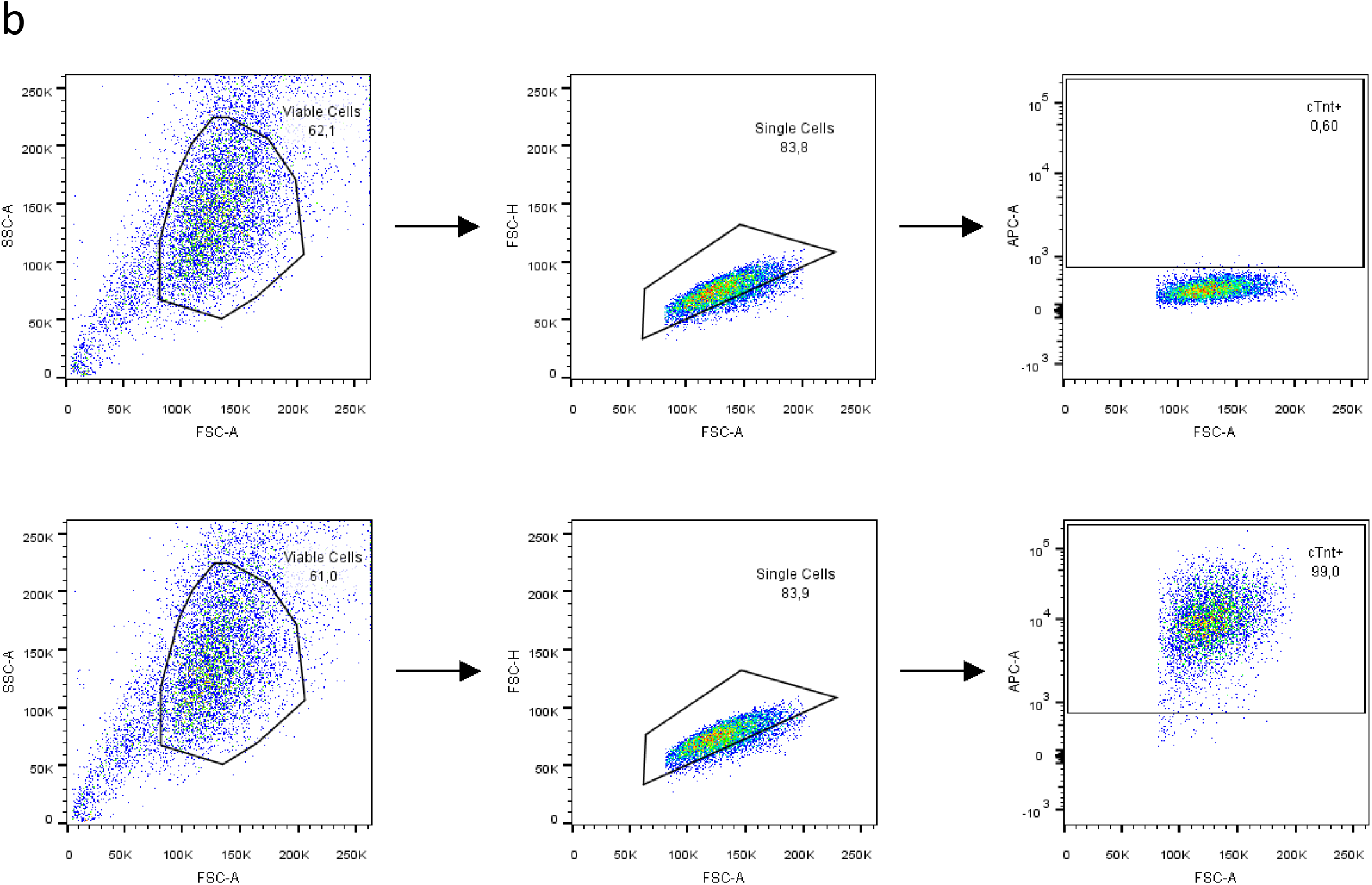
FACS gating strategy example. For all experiments FSC-A vs. SSC-A gates of the starting cell population were used to identify viable cells. Single cells were identified using FSC-A vs. FSC-H gating. Positive populations were determined by the specific antibodies or native EYFP/EGFP signal, which were distinct from negative populations. Isotype control or unedited cells were used to distinguish between background and marker-positive events.

**Supplementary Figure 6:**
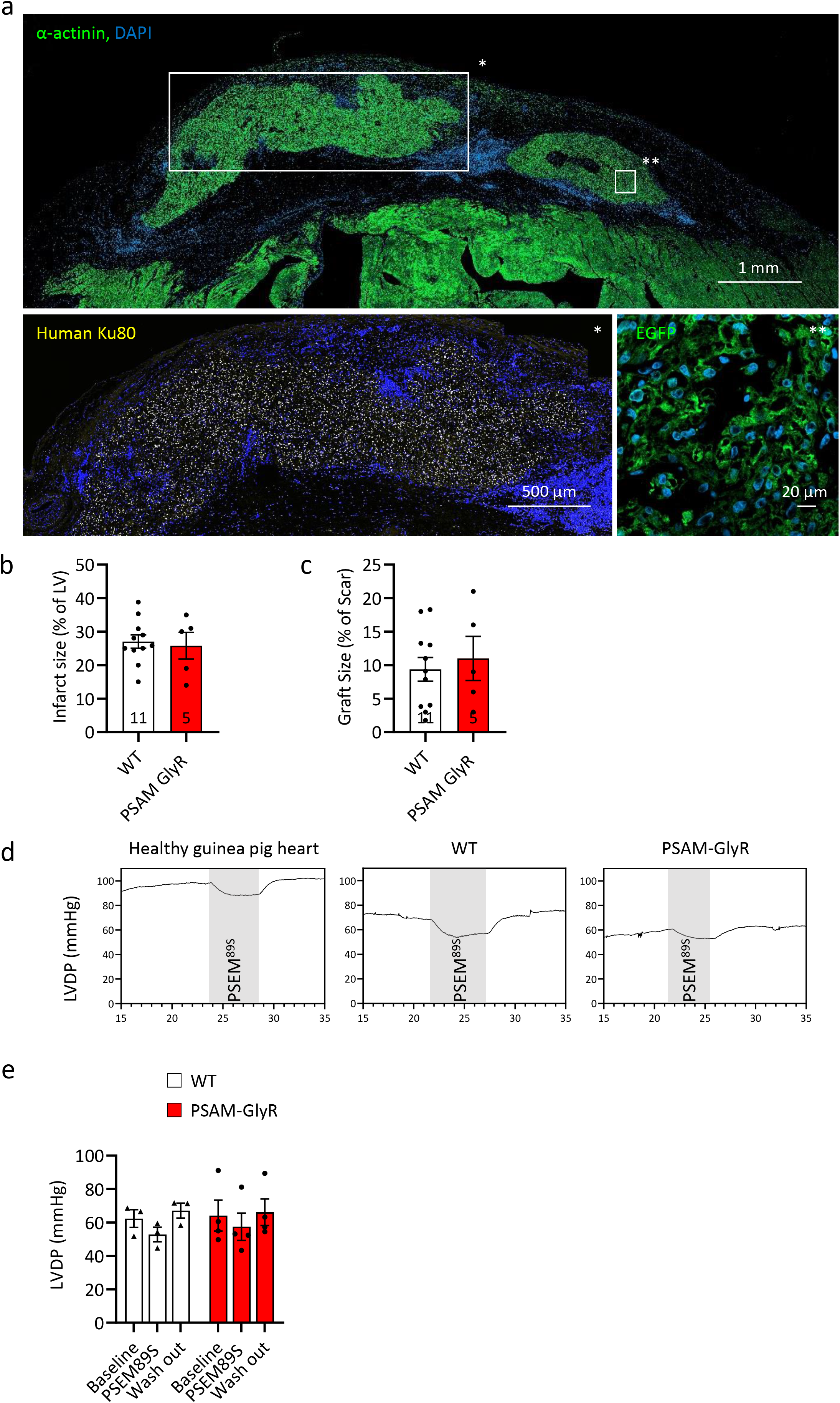
PSAM-GlyR cardiomyocyte transplantation study. **a)** Histology of the anterior wall in short axis stained for α-actinin (low magnification) and human Ku80 and EGFP in high magnification 4 weeks after PSAM-GlyR cardiomyocyte transplantation. **b** and **c**) Histological quantifications of infarct (**b**) and graft size (**c**). **d**) Left ventricular pressure measurement of i) one healthy guinea pig heart, ii) one injured heart four weeks after WT-cardiomyocyte injection and iii) one injured heart four weeks after PSAM-GlyR cardiomyocyte transplantation in the Langendorff-system. Ten beats were averaged for each data point. PSEM^89S^ was given over a time period of 5 minutes. **e**) Quantification of LVDP of Langendorff-perfused hearts four weeks after transplantation of PSAM-GlyR or WT-cardiomyocytes respectively. Mean ± SEM are shown, each data point represents one heart. LVDP: Left ventricular developed pressure.

**Supplementary Figure 8:**
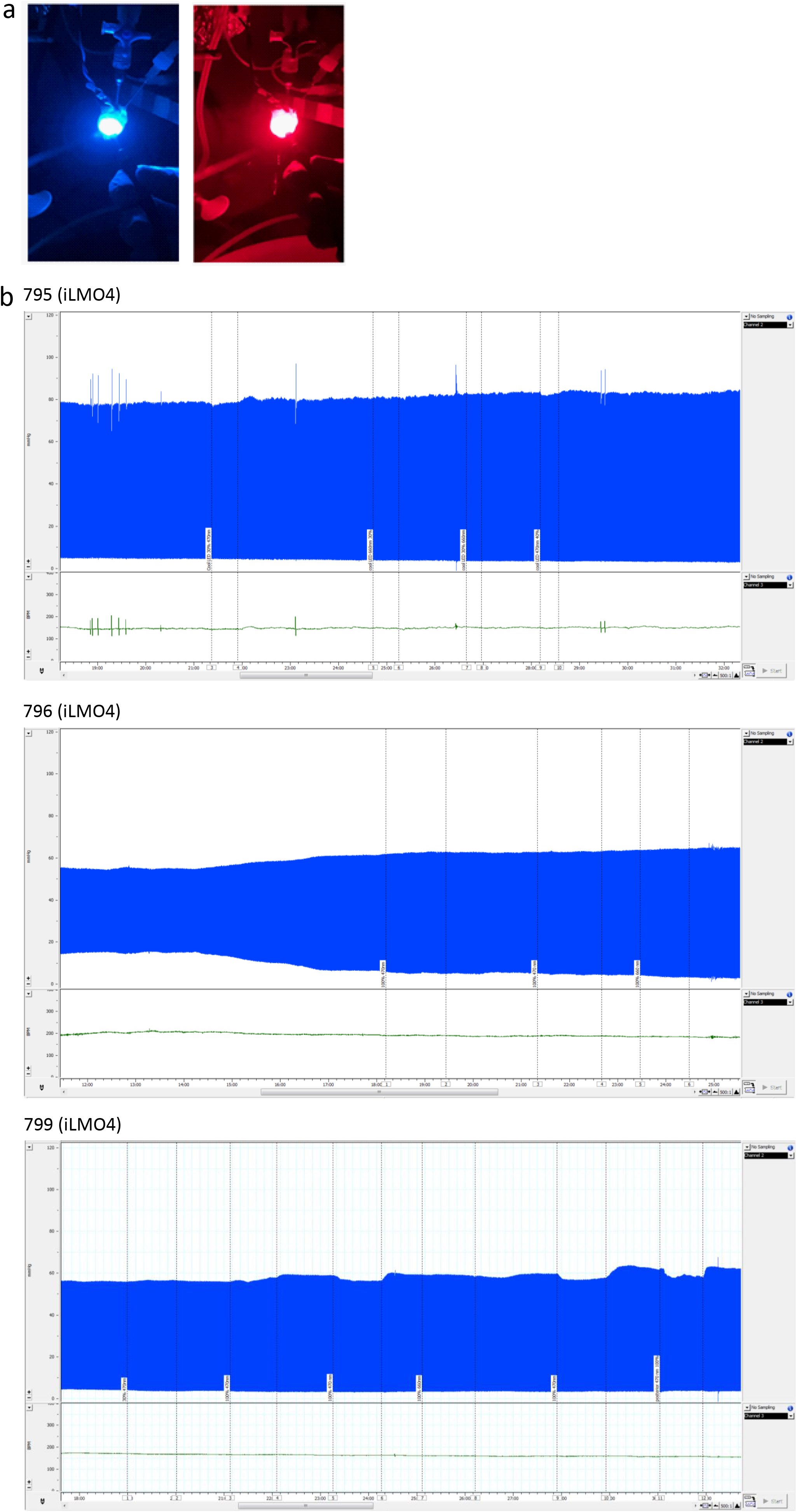
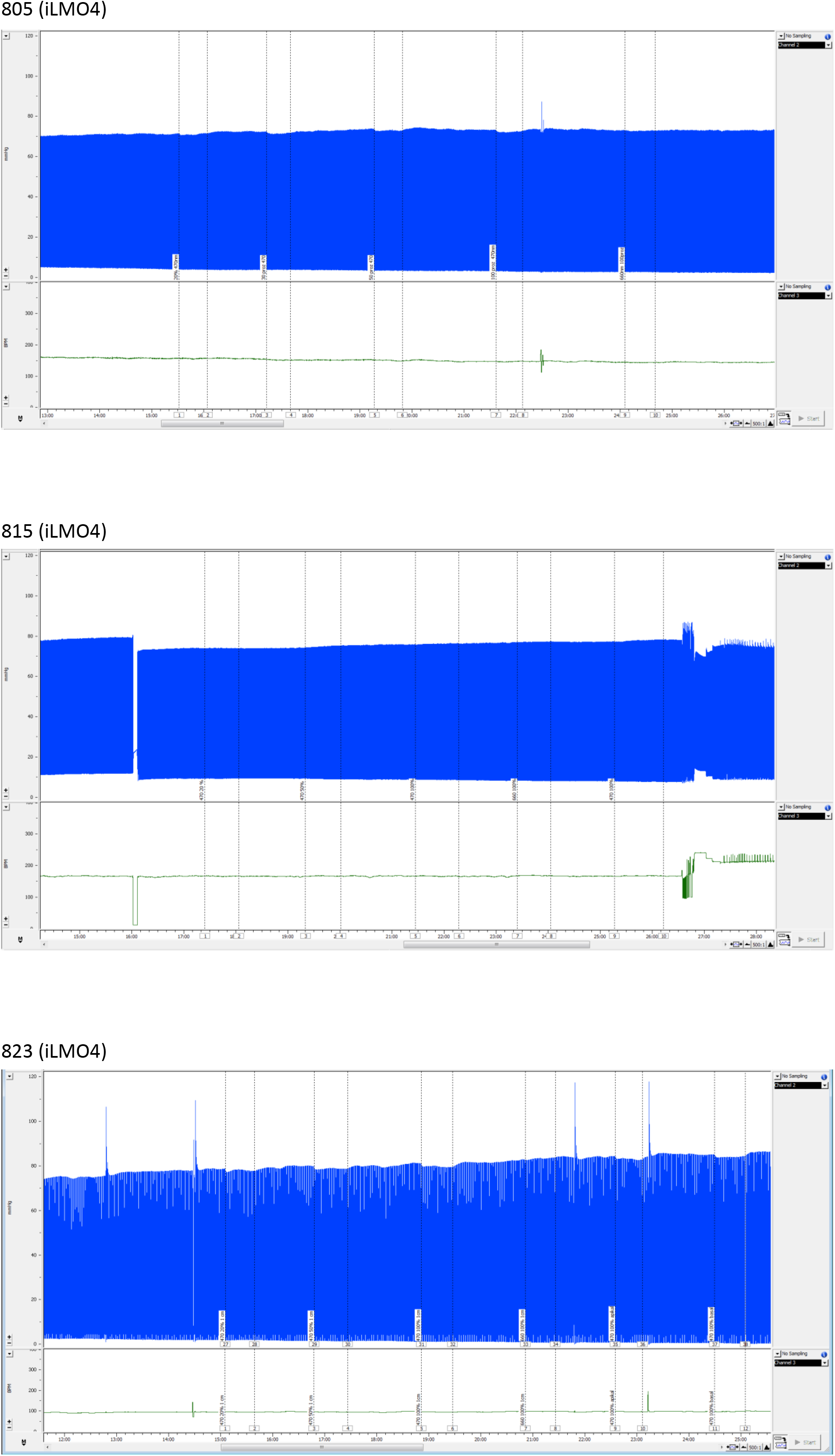
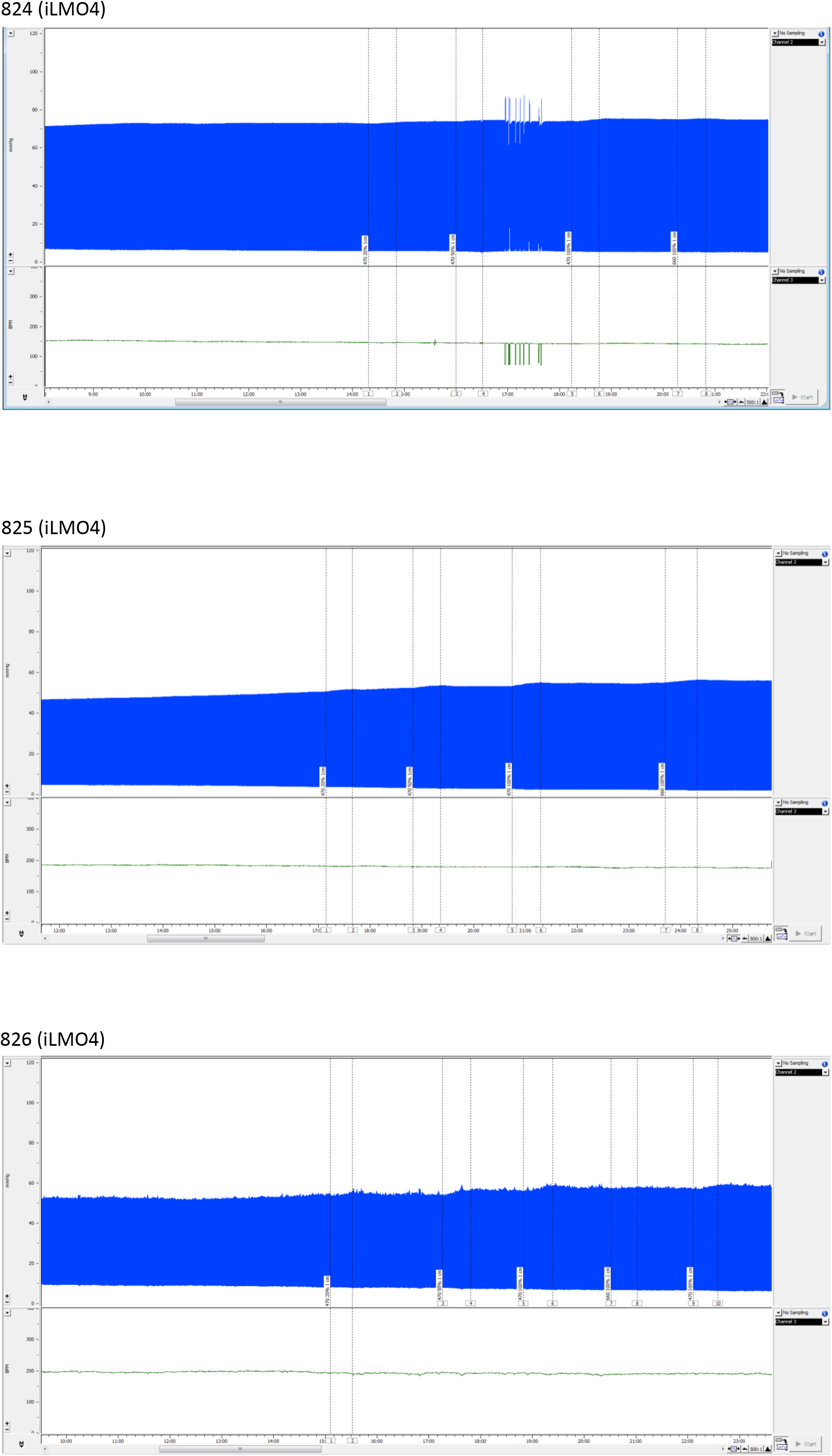
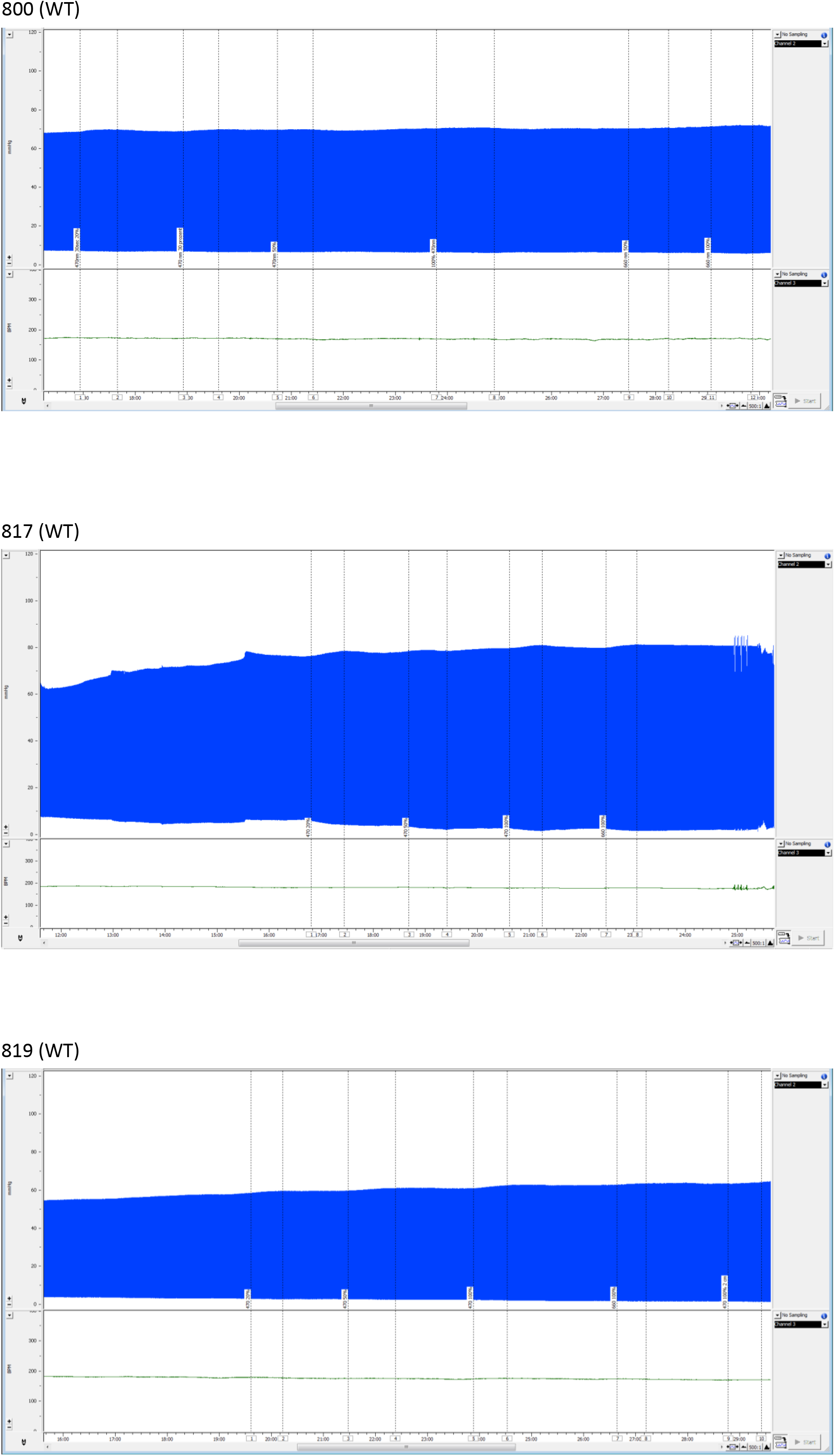
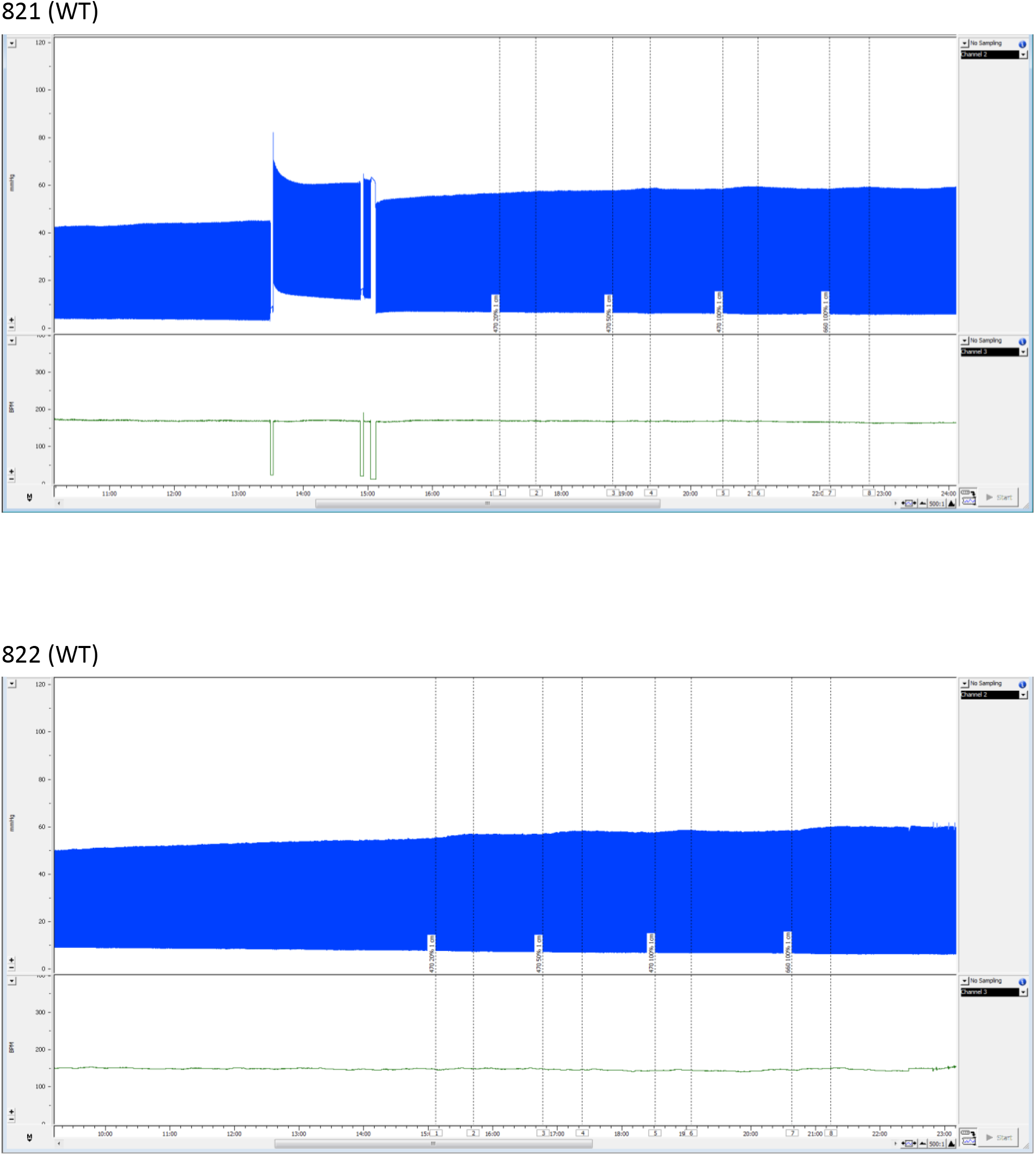
Langendorff-perfusion of iLMO4 and WT hearts. **a**) Photographs of Langendorff-perfused hearts during photostimulation with blue and red light, respectively. **b**) Original LV pressure-recordings from all iLMO4 and WT Langendorff-perfused hearts.

## References

1. Querdel, E. et al. Human Engineered Heart Tissue Patches Remuscularize the Injured Heart in a Dose-Dependent Manner. Circulation 143, 1991–2006 (2021).

2. Laflamme, M. a et al. Cardiomyocytes derived from human embryonic stem cells in pro-survival factors enhance function of infarcted rat hearts. Nat. Biotechnol. 25, 1015–24 (2007).

3. Liu, Y.-W. W. et al. Human embryonic stem cell–derived cardiomyocytes restore function in infarcted hearts of non-human primates. Nat. Biotechnol. 36, 597–605 (2018).

4. Romagnuolo, R. et al. Human Embryonic Stem Cell-Derived Cardiomyocytes Regenerate the Infarcted Pig Heart but Induce Ventricular Tachyarrhythmias. Stem Cell Reports 12, 967–981 (2019).

5. Cyranoski, D. ‘Reprogrammed’ stem cells approved to mend human hearts for the first time. Nature 557, 619–620 (2018).

6. Eschenhagen, T. et al. Cardiomyocyte regeneration: A consensus statement. Circulation 136, 680–686 (2017).

7. Roth, G. A. A. et al. Global Burden of Cardiovascular Diseases and Risk Factors, 1990-2019: Update From the GBD 2019 Study. J. Am. Coll. Cardiol. 76, 2982–3021 (2020).

8. Velagaleti, R. S. et al. Long-term trends in the incidence of heart failure after myocardial infarction. Circulation 118, 2057–62 (2008).

9. Weinberger, F. & Eschenhagen, T. Cardiac Regeneration: New Hope for an Old Dream. Annu. Rev. Physiol. 83, annurev-physiol-031120-103629 (2021).

10. Weinberger, F. et al. Cardiac repair in guinea pigs with human engineered heart tissue from induced pluripotent stem cells. Sci. Transl. Med. 8, 363ra148 (2016).

11. Zhao, M. et al. Cyclin d2 overexpression enhances the efficacy of human induced pluripotent stem cell-derived cardiomyocytes for myocardial repair in a swine model of myocardial infarction. Circulation 144, 210–228 (2020).

12. Zhu, K. et al. Lack of Remuscularization Following Transplantation of Human Embryonic Stem Cell-Derived Cardiovascular Progenitor Cells in Infarcted Nonhuman Primates. Circ. Res. 122, 958–969 (2018).

13. Magnus, C. J. et al. Chemical and genetic engineering of selective ion channel-ligand interactions. Science (80-.). 333, 1292–1296 (2011).

14. Wietek, J. et al. Conversion of Channelrhodopsin into a Light-Gated Chloride Channel. Science (80-.). 344, 409–412 (2014).

15. Park, S. Y. et al. Novel luciferase-opsin combinations for improved luminopsins. J. Neurosci. Res. (2017) doi:10.1002/jnr.24152.

16. Wietek, J. et al. An Imporved Chloride-Conducting Channelrhodopsin for Light-Induced Inhibition of Neuronal Activity In Vivo. Sci. Rep. 5, 14807 (2015).

17. Qian, K. et al. A simple and efficient system for regulating gene expression in human pluripotent stem cells and derivatives. Stem Cells 32, 1230–1238 (2014).

18. Breckwoldt, K. et al. Differentiation of cardiomyocytes and generation of human engineered heart tissue. Nat. Protoc. 12, 1177–1197 (2017).

19. Guo, L., Dong, Z. & Guthrie, H. Validation of a guinea pig Langendorff heart model for assessing potential cardiovascular liability of drug candidates. J. Pharmacol. Toxicol. Methods 60, 130–151 (2009).

20. Zacchigna, S. et al. Towards standardization of echocardiography for the evaluation of left ventricular function in adult rodents: A position paper of the ESC Working Group on Myocardial Function. Cardiovascular Research vol. 117 43–59 (2021).

21. Shiba, Y. et al. Human ES-cell-derived cardiomyocytes electrically couple and suppress arrhythmias in injured hearts. Nature 489, 322–325 (2012).

22. Liao, R., Podesser, B. K. & Lim, C. C. The continuing evolution of the Langendorff and ejecting murine heart: new advances in cardiac phenotyping. Am. J. Physiol. Heart Circ. Physiol. 303, H156-67 (2012).

23. Chong, J. J. H. et al. Human embryonic-stem-cell-derived cardiomyocytes regenerate non-human primate hearts. Nature 510, 273–277 (2014).

24. Laflamme, M. A. et al. Formation of human myocardium in the rat heart from human embryonic stem cells. Am. J. Pathol. 167, 663–671 (2005).

25. Schaaf, S. et al. Human Engineered Heart Tissue as a Versatile Tool in Basic Research and Preclinical Toxicology. PLoS One 6, e26397 (2011).

26. Lemme, M. et al. Chronic intermittent tachypacing by an optogenetic approach induces arrhythmia vulnerability in human engineered heart tissue. Cardiovasc. Res. 116, 1487–1499 (2020).

